# Efficient transfection of Atlantic salmon primary hepatocyte cells for functional assays and gene editing

**DOI:** 10.1101/2022.10.20.513028

**Authors:** Alex K. Datsomor, Ragnhild Wilberg, Jacob S. Torgersen, Simen R. Sandve, Thomas N. Harvey

## Abstract

The expansion of genomic resources for Atlantic salmon over the past half decade has enabled efficient interrogation of genetic traits by large-scale correlation of genotype to phenotype. Moving from correlation to causation will require genotype-phenotype relationships to be tested experimentally in a cost-efficient and cell context relevant manner. To enable such future experiments, we have developed a method for the isolation and genetic manipulation of primary hepatocytes from Atlantic salmon for use in heterologous expression, reporter assay, and gene editing experiments. We chose the liver as the tissue of interest because it is the metabolic hub and many current Atlantic salmon research projects focus on understanding metabolic processes to improve traits such as growth rate, total fat content, and omega-3 content. We find that isolated primary hepatocytes are optimally transfected with both plasmid and ribonucleoprotein using a Neon electroporator at 1400 V, 10 ms, and 2 pulses. Transfection efficiency with plasmid and cutting efficiency with ribonucleoprotein was optimally 46% and 60%, respectively. We also demonstrate a 26-fold increase in luciferase expression under the promoter of the key liver metabolic gene, *elovl5b*, compared to empty vector, in line with expected liver-specific expression. Taken together, this work provides a valuable resource enabling transfection and gene editing experiments in a context relevant and cost-effective system.

## 1. Introduction

The release of a high-quality reference genome for Atlantic salmon in 2016 (Lien et al. 2016) enabled efficient scans for genome wide genotype-phenotype associations. This resulted in more efficient breeding programs through marker assisted- and genomic selection (Houston et al. 2020) and a step-change in our ability to understand the genetics of traits in domestic and wild populations (Barson et al. 2015). Yet, the majority of genotype-trait associations is a result of linkage disequilibrium to unknown causative variants, and this limits the utility of such associations in wild population management and breeding (Daetwyler et al. 2013).

Moving forward, development of genomic resources and tools to help tease apart correlation from causation will be of great importance for applied and basic research on the genetics of Atlantic salmon traits. One such initiative is Functional Annotation of ANimal Genomes (FAANG) (Andersson et al. 2015; Clark et al. 2020), which aims to systematically generate and archive functional genomics phenotypes such as gene expression, chromatin accessibility, and histone tail modifications across tissues and developmental stages, and cell types. This data can then be used to provide genome regulatory context of genetic variants associated with phenotypes, which takes us one step closer to causal relationships. However, to fully bridge the genotype-phenotype gap, and dissect out gene function and causal variants using reporter assays or CRISPR-based approaches, efficient protocols for transfection of DNA or proteins into cells are needed.

Several studies have demonstrated transfection of Atlantic salmon cell lines using electroporation of DNA and CRISPR components (Gratacap et al. 2020; Schiøtz et al. 2011), with reports of transfection efficiency ranging between 10-90% (Schiøtz et al. 2011). Even though continuous cell lines can be excellent systems to study many aspects of cell biology and genetics, the immortalization process and lab-evolution often lead to cells with different properties than their tissue of origin (Ben-David et al. 2018; Lopes-Ramos et al. 2017). Hence, for certain applications primary cell cultures are preferred over continuous cell lines, as they often bear stronger functional resemblance to the cells *in vivo* (Zeilinger et al. 2016; Nagarajan et al. 2019). Unfortunately, for Atlantic salmon primary cells efficient transfection protocols are lacking. Standard chemical transfection has proven extremely inefficient in primary cells of Atlantic salmon and other teleosts. For example, chemical transfection of primary gill and liver cells from rainbow trout and Atlantic salmon, respectively, failed to reach 1% transfection efficiency (Wilberg 2020; Romoren et al. 2005). Furthermore, only a few cell lines from Atlantic salmon are in fact available for the research community, and none from key metabolic tissues like the liver. There is therefore a pressing need to develop efficient transfection protocols which will enable functional genomics in primary cells, as well as aid in developing new Atlantic salmon cell lines. The current study describes an efficient transfection protocol for Atlantic salmon primary hepatocytes and employs the optimal protocol for functional assays and CRISPR/Cas9 based studies.

## 2. Materials and Methods

### 2.1. Isolation of Atlantic salmon primary hepatocytes

Atlantic salmon (*Salmo salar*) parrs of 100-400 g were obtained from the Centre for Fish Research, NMBU. Fish were euthanized by a sharp blow to the head, and the liver immediately perfused for 10 minutes with ice-cold wash buffer, pH=7.4 (1× Hank’s Balanced salt solution, HBSS without Mg^2+^/Ca^2+^, 1 mM EDTA, 10 mM HEPES) via the portal vein. Liver was subsequently perfused for 10 minutes with ice-cold collagenase buffer, pH=7.5 (1× HBSS with Mg^2+^/Ca^2+^, 10 mM HEPES, 150 U/ml collagenase) via the portal vein, gently torn into small pieces and incubated for 1 hr in sterile Erlenmeyer flask with collagenase buffer at 15 °C under atmospheric conditions with continuous slow stirring on a magnetic stirrer. Dissociated hepatocytes were thereafter filtered through a 100 μm cell strainer and rinsed with ice-cold Leibovitz’s L-15 medium with GlutaMAX^™^ supplement (ThermoFisher Scientific). Hepatocytes were harvested at 100× g for 5 min at 4 °C, resuspended in 5 ml 1x HBSS (without Mg^2+^/Ca^2+^) and spun down again at 100× g at 4 °C for 5 mins. Then, the cells were resuspended in 5 ml 1× HBSS (without Mg^2+^/Ca^2+^) and counted using the hemocytometer with trypan blue. Polyethylenimine coated plates were used to facilitate attachment of hepatocytes in all experiments. Protocol has been published on protocols.io (Datsomor et al. 2022).

### 2.2. Optimization of electroporation-based transfection protocol

Electroporation of salmon primary hepatocytes was performed using the neon® transfection system (Invitrogen) in accordance with the manufacturer’s protocol. Approximately 4×10^5^ hepatocytes were transfected with 3 μg of reporter plasmid, pEGFP-N1-FLAG (Addgene# 60360) per well of a 6-well plate. Transfection was performed using electroporation programs with varying voltage, pulse width and pulse number (Table 1). Transfected cells were incubated in L15 medium (L15 GlutaMAX^™^, 5% fetal bovine serum, without antibiotics) at 15 °C under atmospheric conditions. 24 hours after transfection, media was replaced with fresh complete L15 medium (L15 GlutaMAX^™^, 5% foetal bovine serum, 1× streptomycin-Penicillin) and cells were incubated at 15 °C under atmospheric conditions for an additional 24 hrs. At 48 hours post-transfection, the impact of the various programs on cell viability was assessed by Resazurin viability assay (Sigma) in accordance with manufacturer’s protocol. Successful transfection was evaluated by GFP expression using the ZEISS fluorescence microscope and the proportion of transfected cells determined by flow cytometry using the Amnis Cellstream (Luminex).

**Table 1:** Neon electroporator parameters for the four programs tested.

| Program ID | Voltage (V) | Pulse width (ms) | Pulse number |
| --- | --- | --- | --- |
| P2 | 1400 | 20 | 1 |
| P9 | 1400 | 30 | 1 |
| P16 | 1400 | 20 | 2 |
| P20 | 1150 | 30 | 2 |

### 2.3. Implementation of optimal transfection protocol for comparative promoter study

To demonstrate the importance of successful transfection and its potential for functional studies in primary hepatocytes, we employed the optimal transfection program in a luciferase assay for the Atlantic salmon *elovl5b* gene that has shown a liver-specific expression pattern (Morais et al. 2009).

#### 2.3.1. Preparation of promoter luciferase construct

We generated a promoter luciferase reporter construct for *elovl5b* using the pGL4.10[luc2] vector (Promega, GenBank accession# AY738222), which contains the firefly luciferase reporter gene. This vector includes the wild-type “full-length” promoter (pGL4.10-elovl5bWT) which contains the *elovl5b* promoter region NC_027327.1: 27244001 – 27245560 (Supplemental data 1). This vector is hereafter termed as elovl5bWT. The promoter was amplified from Atlantic salmon genomic DNA using the Platinum^™^ SuperFi PCR Master mix (ThermoFisher) with primers containing 15 bp tails homologous to cloning sites within the pGL4.10[luc2] vector which is necessary for cloning using the In-Fusion® HD cloning kit (Takara). Primer sequences are indicated in Table S1. The promoter reporter vector was isolated using the ZymoPure plasmid miniprep kit and confirmed by Sanger sequencing (LightRun Tube Sequencing Service, Eurofins).

#### 2.3.2. Transfection of promoter luciferase constructs and luciferase assay

Approximately 1.0 - 1.5 ×10^5^ isolated primary hepatocytes were co-transfected per well in 24-well plates with 1.5 μg of promoter reporter construct and 0.5 μg of the reference reporter construct, pGL4.75[hRluc/CMV] (Promega), encoding Renilla luciferase. Transfection was performed using electroporation program P16 (Table 1). At 24 hours post-transfection, fresh complete L15 medium (L15 GlutaMAX™, 5% foetal bovine serum, 1× streptomycin-Penicillin) was added to transfected cells and incubated at 15 °C under atmospheric conditions for an additional 24 hours. To quantify firefly and Renilla luciferase activities, medium on cells was replaced with 100 μl of Dulbecco’s Modified Eagle’s medium (Sigma) and 100 μl Dual-Glo® luciferase reagent (Promega) per well and incubated for approximately 30 min. Luminescence were read on Synergy H1 Hybrid multi-mode microplate reader (Bio Tek). Luminescence from Renilla luciferase activities was measured 10 min after adding 100 μl of Dual-Glo® Stop & Glo® reagent. Firefly luminescence was normalized to Renilla luciferase luminescence.

### 2.4. Implementation of optimal transfection protocol for genome-editing using RNPs

To identify the optimal program for genome editing using ribonucleoproteins (RNPs) we designed a guide RNA to Atlantic salmon tp53 (NCBI geneID: 106602901) and performed RNP electroporation using the same programs in table 1 as described above.

#### 2.4.1. Preparation and electroporation of RNPs

RNP complexes were prepared according to the Alt-R CRISPR-Cas9 system protocol from IDT. In brief, crRNA:tracrRNA duplexes were made by diluting 2.2 μl of Ssal_tp53_crRNA (200 μM) and 2.2 μl of tracrRNA (200 μM) in 5.6 μl of IDTE buffer. Cas9 protein was prepared by mixing 12 μl of Alt-R Cas9 protein (62 μM) with 8 μl of buffer R (Invitrogen). gRNA duplexes and Cas9 were then mixed gently and incubated at room temperature for 10 minutes. 1 μl of prepared RNPs was added to 9 μl of cells resuspended in buffer R just before electroporation. After electroporation, cells were distributed directly into wells of a 24 well-plate containing 1 mL of culture medium (L15, 5% FBS) without antibiotics. After 24 hours at 15°C, media was replaced with culture medium containing antibiotics and antimycotics (L15, 5% FBS, 100 U/mL PenStrep, 2.5 μg/mL Amphotericin B) and placed at 15°C. Cells were harvested 72 hours later by washing twice with PBS (pH 7.2) and incubating with 100 μl 0.25% trypsin/EDTA (Invitrogen) until cells detached. 400 μl of culture medium containing FBS was then added to inactivate the trypsin. Cells were centrifuged at 150× g for 5 minutes and washed twice in PBS, then stored at −20 °C.

#### 2.4.2. Determination of CRISPR-editing efficiency

Genomic DNA was extracted from frozen cell pellets according to the QIAGEN blood and tissue kit protocol. Genomic loci containing the Cas9 cut site were amplified by PCR using the primers Ssal_tp53_seq_fwd and Ssal_tp53_seq_rev (Table 2). PCR reactions were set up as follows: 25 μl Platinum II hot start green PCR master mix (Invitrogen), 1 μl each primer (10 μM), 7.5 μl gDNA (~10 ng/μl), 15.5 μl nuclease free H_2_O. Thermocycler conditions were as follows: Initial denaturation of 94 °C for 2 minutes followed by 35 cycles of 94 °C denaturation for 15 seconds, 60 °C annealing for 15 seconds, 68 °C extension for 15 seconds. A single clear band at 844 bp was obtained for all reactions. PCR reactions were cleaned up using a QIAquick PCR cleanup kit (QIAGEN) and Sanger sequencing of PCR products using Ssal_tp53_seq_rev was performed by Eurofins genomics. Cas9 cutting efficiency was determined by ICE deconvolution (Conant et al. 2022) of sanger sequencing traces using default parameters.

**Table 2:**
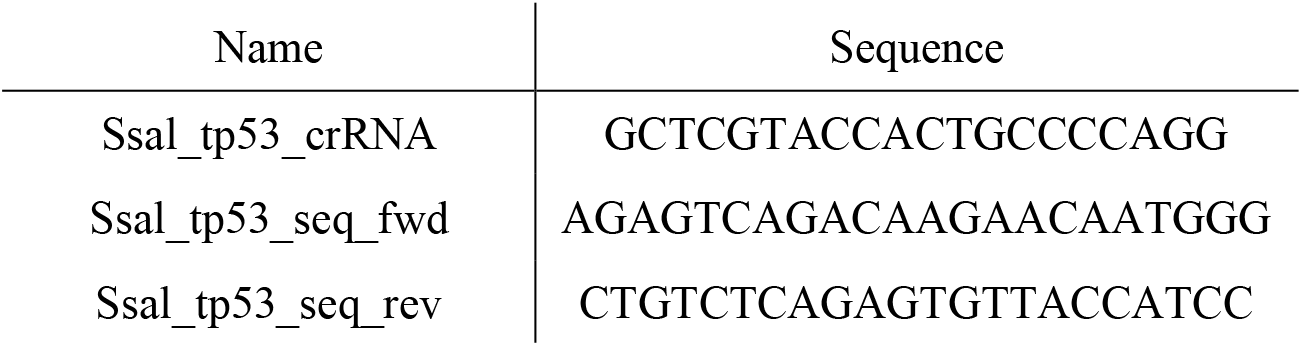
tp53 targeting crRNA sequence and primer sequences for PCR and sequencing of targeted locus.

### 2.5 Statistical analysis

The effects of different programs on transfection efficiency and viability were determined by one-way analysis of variance (ANOVA), followed by Tukey’s multiple comparison test with a p-value cutoff of 0.05. Comparison of luminescence from different promoters was assessed using a student’s t-test and a p-value cutoff of 0.05. Results shown are representative of 3 independent experiments. All statistical analysis was performed in R version 4.2.1. (Team 2020) using RStudio (Team 2019)

## 3. Results

### 3.1 Isolation of primary hepatocytes

We isolated primary hepatocytes from freshwater stage Atlantic salmon (100-400 g) by perfusing the liver *in situ* through the portal vein with wash buffer to remove erythrocytes, followed by collagenase to digest the extracellular matrix, and several filtration and washing steps to obtain a suspension of single cells (Figure 1). The isolated primary hepatocytes grew optimally at 15 °C under atmospheric condition in L15 medium supplemented with 5% FBS. We found that a liver from a 200 g fish would typically yield between 2×10^7^ and 4×10^7^ cells with a viability of 80-95% prior to electroporation, as determined by trypan blue staining and counting with a hemocytometer. The isolated primary hepatocytes were mostly dispersed single cells and spherical in shape. We also observed some oval shaped erythrocytes, however these were greatly reduced by *in situ* perfusion. Isolated hepatocytes did not attach to ordinary non-coated culture plates, but when growth surfaces were coated with polyethylenimine (branched) the cells attached optimally at a density of 7-10 ×10^4^ cells/cm^2^. 24 to 48 hours after attachment cells formed flattened aggregates that would slowly expand. Cells remained viable under our conditions for at least three weeks.

**Figure 1:**
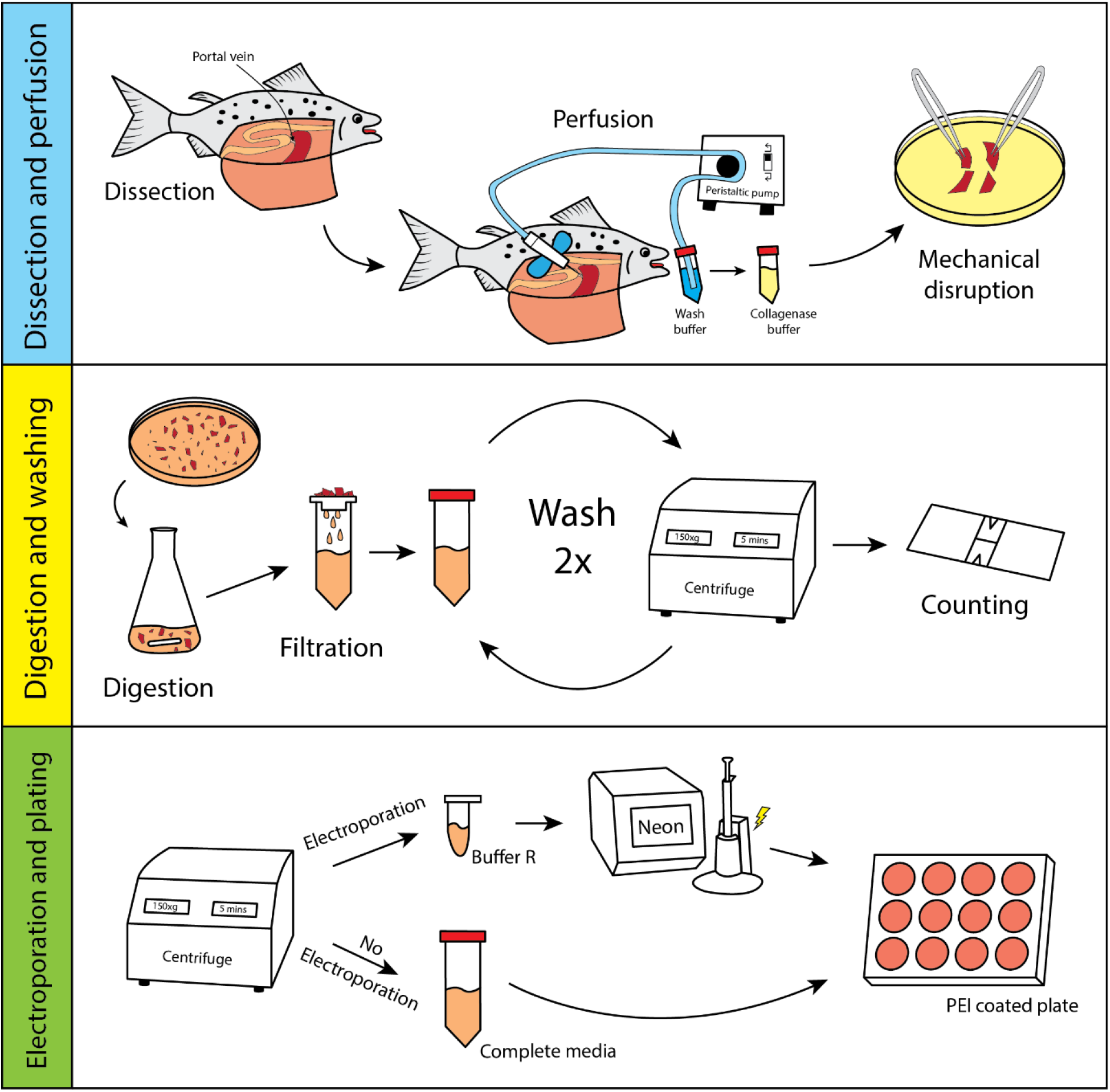
Schematic of the primary hepatocyte isolation procedure and electroporation using the Neon electroporation system.

### 3.2. Transfection of GFP expression plasmid and functional evaluation of *elovl5b* promoter

We tested 24 electroporation conditions using pGL4.75 which encodes for Renilla luciferase under the control of the CMV promoter. We found a positive correlation between voltage and transfection efficiency, with 1400 V, 2 pulses having the highest efficiency (Figure S1). Of these conditions we selected four, three high efficiency and one low efficiency, for further analysis by transfecting with a plasmid encoding GFP under the control of the CMV promoter. Transfection efficiency was measured by fluorescent microscopy and flow cytometry. We found P16 (1400 V, 20 ms, 2 pulses) to have the highest transfection efficiency of 46% (Figure 2A, B). P2, P9, and P20 had electroporation efficiencies of 9%, 33%, and 36%, respectively. Cell viability, as measured by the conversion of resazurin to resorufin, decreased after electroporation for all conditions with no clear differences between the four conditions (Figure 2C).

**Figure 2:**
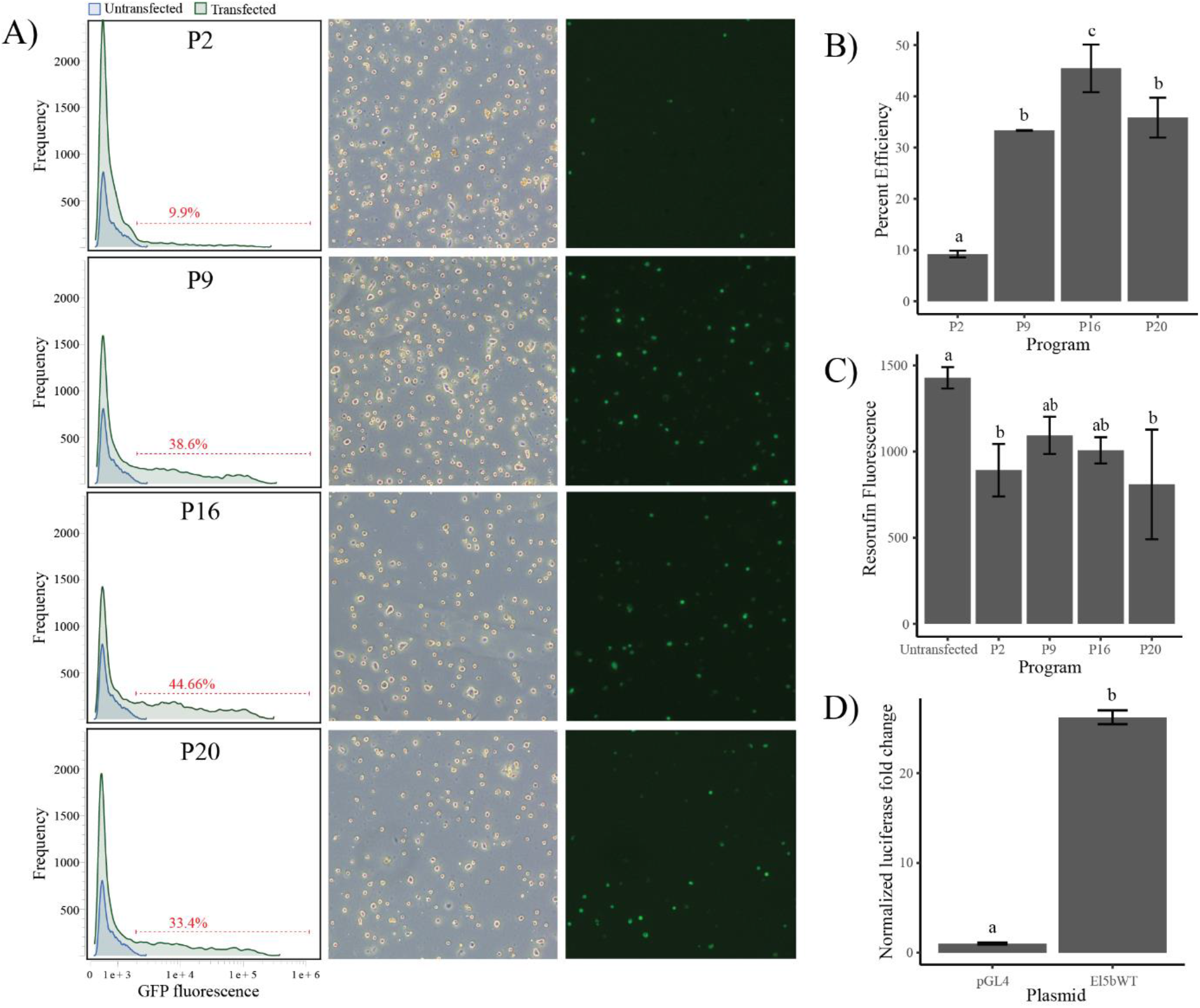
Impact of electroporation programs on transfection efficiency and viability and demonstration of functional reporter assay in Atlantic salmon primary hepatocytes. A and B) The different electroporation programs assessed show varying effects on transfection of hepatocytes with program P16 (Table 1) giving highest transfection efficiency. Transfection efficiency was measured by flow cytometry. C) Evaluation of cell viability by resazurin viability assay at 48-hr post transfection showed slight reduction in cell viability for all programs compared to non-transfected control. D) Normalized luciferase signal for Atlantic salmon *elovl5b* promoter compared to the empty vector.

To showcase the utility of the transfection protocol for primary liver cells we performed a luciferase promoter-reporter assay using the promoter of a known liver centric gene involved in the fatty acid metabolism (*elovl5b*). The *elovl5b* promoter showed significant (p< 0.05) 26-fold increase in luciferase signal compared to the empty vector (Figure 2D).

### 3.3 CRISPR Cas9 gene editing

To test the effectiveness of gene editing by RNP electroporation in Atlantic salmon primary cells we designed a single guide RNA (sgRNA) to one of the three salmon P53 genes (NCBI geneID:106602901). We then combined this with Cas9 protein to form RNPs and electroporated using the four conditions described above. We found that P16 had the highest cutting efficiency of 60%, P9 and P20 had slightly lower efficiencies of 49% and 54%, respectively, and P2 had the lowest of 31% (Figure 3). ICE deconvolution of sanger sequencing traces showed that the majority of indels were one or two base pair deletions and none were insertions (Figure 3).

**Figure 3:**
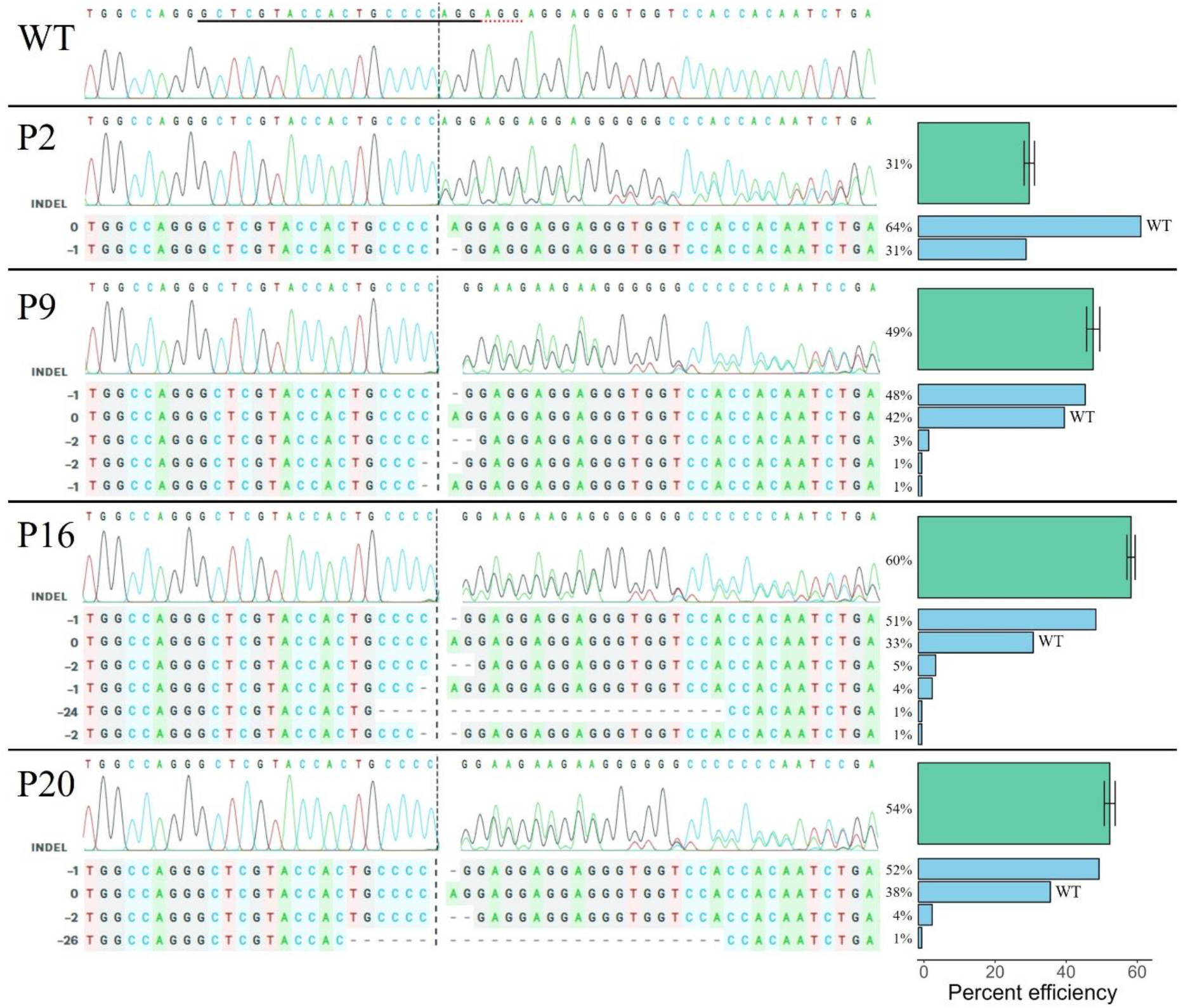
Cutting efficiency of RNPs targeting Atlantic salmon P53 (NCBI geneID:106602901) using four electroporation programs. Sanger sequencing traces for each electroporation program are shown. The gRNA binding site is underlined in the WT trace and the cut site is indicated by a vertical dashed line in all traces. ICE deconvolutions are shown for each program. Bar plots (right) indicate total percent cutting efficiency (green) and contribution of each indel (blue). Indels of length 0 are unaltered WT sequence and negative indels are deletions. No insertions were detected.

## 4. Discussion

High quality *in vitro* experimental cell model systems for commercially important aquaculture species like Atlantic salmon are very important for implementation of modern molecular techniques. The establishment of high-throughput and robust methods at the cutting edge of molecular biology is necessary for advanced research into genetic mechanisms underlying physiological processes. However, no robust protocols for transfection and genetic engineering of primary cells exist in Atlantic salmon. To this end, we have established an efficient transfection protocol for Atlantic salmon primary hepatocytes, the hub of fat and energy metabolism, and demonstrate our ability to employ these cells for functional and CRISPR/Cas9 based studies.

Transfection efficiency of continuous cell lines and primary cells is highly dependent on the cell type and the method used. In our study we measured electroporation efficiency for both plasmid DNA and RNPs and found optimally 46% and 60% efficiencies, respectively. Electroporation of Atlantic salmon TO cells and Atlantic salmon kidney (ASK) cells has achieved a plasmid transfection efficiency up to 90% and 50%, respectively (Schiøtz et al. 2011). Neon electroporation has been shown to efficiently deliver RNP complexes to salmon head kidney (SHK-1) and ASK cells achieving editing efficiencies as high as 90% (Gratacap et al. 2020). This higher efficiency in cell lines compared to our cells is expected because primary cells are notably more difficult to transfect than cell lines.

To our knowledge, our study represents the first protocol of transfection of primary hepatocytes in Atlantic salmon, however many studies have been conducted in mammalian systems. Human umbilical vein endothelial cells have been demonstrated to achieve plasmid electroporation efficiencies of up to 90% with viability greater than 70% (Gresch and Altrogge 2012). Primary hepatocytes typically have lower electroporation efficiencies between 25-40% for plasmid (Chen et al. 2005; Gao et al. 2012) and 52-78% for RNPs (Rathbone et al. 2022) which is more in line with our observations in Atlantic salmon hepatocytes. Interestingly, Chen et al. (2005) was able to double electroporation efficiency in primary mouse hepatocytes by electroporating cells 24 hours after partial hepatectomy (Chen et al. 2005). This demonstrates that cell growth rate is likely a major factor limiting electroporation efficiencies in primary hepatocytes, so increasing growth rate through optimization of growth conditions could be a route to improving transfection efficiency.

High efficiency transfection of primary hepatocytes opens opportunities for modern high-throughput molecular techniques in Atlantic salmon within a physiologically relevant context. For example, massively parallel reporter assays which would enable systematic genome-wide identification of cis-regulatory elements (Wang et al. 2018; Kircher et al. 2019; Klein et al. 2020). The transfection efficiency we obtained in our study is in the order of what is required for these assays as it enables manageable cell numbers and ensures cost-efficient experimental design. In addition, the promoter of *elovl5b* showed a 26-fold increase in activity in the primary hepatocytes consistent with the liver-specific expression pattern of salmon *elovl5b* (Morais et al. 2009), which underscores the physiological semblance between isolated hepatocytes and the liver. Our high cutting efficiency of RNP electroporation in primary hepatocytes will enable metabolically relevant *ex vivo* gene knockout studies in Atlantic salmon. For example, recent studies have knocked out key lipid metabolism genes in Atlantic salmon to study the function *in vivo* (Datsomor et al. 2019a; Datsomor et al. 2019b), however these fish trials are extremely time consuming and costly. *Ex vivo* gene editing of primary hepatocytes would enable quicker turnaround times and allow for the elucidation of a wider range of gene functions. Taken together, our protocol for efficient plasmid transfection and gene editing in primary hepatocytes will open a wide variety of opportunities to study hepatic function in Atlantic salmon.

## References

Andersson, L., A.L. Archibald, C.D. Bottema, R. Brauning, S.C. Burgess et al., 2015 Coordinated international action to accelerate genome-to-phenome with FAANG, the Functional Annotation of Animal Genomes project. Genome Biology 16 (1).

Barson, N.J., T. Aykanat, K. Hindar, M. Baranski, G.H. Bolstad et al., 2015 Sex-dependent dominance at a single locus maintains variation in age at maturity in salmon. Nature 528 (7582):405–408.

Ben-David, U., B. Siranosian, G. Ha, H. Tang, Y. Oren et al., 2018 Genetic and transcriptional evolution alters cancer cell line drug response. Nature 560 (7718):325–330.

Chen, N.K., J. Sivalingam, S.Y. Tan, and O.L. Kon, 2005 Plasmid-electroporated primary hepatocytes acquire quasi-physiological secretion of human insulin and restore euglycemia in diabetic mice. Gene Ther 12 (8):655–667.

Clark, E.L., A.L. Archibald, H.D. Daetwyler, M.A.M. Groenen, P.W. Harrison et al., 2020 From FAANG to fork: application of highly annotated genomes to improve farmed animal production. Genome Biology 21 (1).

Conant, D., T. Hsiau, N. Rossi, J. Oki, T. Maures et al., 2022 Inference of CRISPR Edits from Sanger Trace Data. CRISPR J 5 (1):123–130.

Daetwyler, H.D., M.P.L. Calus, R. Pong-Wong, G. De Los Campos, and J.M. Hickey, 2013 Genomic Prediction in Animals and Plants: Simulation of Data, Validation, Reporting, and Benchmarking. Genetics 193 (2):347–365.

Datsomor, A.K., R.E. Olsen, N. Zic, A. Madaro, A.M. Bones et al., 2019a CRISPR/Cas9-mediated editing of Δ5 and Δ6 desaturases impairs Δ8-desaturation and docosahexaenoic acid synthesis in Atlantic salmon (Salmo salar L.). Scientific Reports 9 (1).

Datsomor, A.K., R. Wilberg, J.S. Torgersen, S.R. Sandve, and T.N. Harvey, 2022 Transfection of Atlantic salmon primary hepatocytes. protocols.io.

Datsomor, A.K., N. Zic, K. Li, R.E. Olsen, Y. Jin et al., 2019b CRISPR/Cas9-mediated ablation of elovl2 in Atlantic salmon (Salmo salar L.) inhibits elongation of polyunsaturated fatty acids and induces Srebp-1 and target genes. Scientific Reports 9 (1).

Gao, S., E. Seker, M. Casali, F. Wang, S.S. Bale et al., 2012 Ex Vivo Gene Delivery to Hepatocytes: Techniques, Challenges, and Underlying Mechanisms. Annals of Biomedical Engineering 40 (9):1851–1861.

Gratacap, R.L., Y.H. Jin, M. Mantsopoulou, and R.D. Houston, 2020 Efficient Genome Editing in Multiple Salmonid Cell Lines Using Ribonucleoprotein Complexes. Marine Biotechnology 22 (5):717–724.

Gresch, O., and L. Altrogge, 2012 Transfection of difficult-to-transfect primary mammalian cells. Methods Mol Biol 801:65–74.

Houston, R.D., T.P. Bean, D.J. Macqueen, M.K. Gundappa, Y.H. Jin et al., 2020 Harnessing genomics to fast-track genetic improvement in aquaculture. Nature Reviews Genetics 21 (7):389–409.

Kircher, M., C. Xiong, B. Martin, M. Schubach, F. Inoue et al., 2019 Saturation mutagenesis of twenty disease-associated regulatory elements at single base-pair resolution. Nature Communications 10 (1).

Klein, J.C., V. Agarwal, F. Inoue, A. Keith, B. Martin et al., 2020 A systematic evaluation of the design and context dependencies of massively parallel reporter assays. Nature Methods 17 (11): 1083–1091.

Lien, S., B.F. Koop, S.R. Sandve, J.R. Miller, M.P. Kent et al., 2016 The Atlantic salmon genome provides insights into rediploidization. Nature 533 (7602):200–205.

Lopes-Ramos, C.M., J.N. Paulson, C.-Y. Chen, M.L. Kuijjer, M. Fagny et al., 2017 Regulatory network changes between cell lines and their tissues of origin. BMC Genomics 18 (1).

Morais, S., O. Monroig, X. Zheng, M.J. Leaver, and D.R. Tocher, 2009 Highly Unsaturated Fatty Acid Synthesis in Atlantic Salmon: Characterization of ELOVL5-and ELOVL2-like Elongases. Marine Biotechnology 11 (5):627–639.

Nagarajan, S.R., M. Paul-Heng, J.R. Krycer, D.J. Fazakerley, A.F. Sharland et al., 2019 Lipid and glucose metabolism in hepatocyte cell lines and primary mouse hepatocytes: a comprehensive resource for in vitro studies of hepatic metabolism. Am J Physiol Endocrinol Metab 316 (4):E578–E589.

Rathbone, T., I. Ates, L. Fernando, E. Addlestone, C.M. Lee et al., 2022 Electroporation-Mediated Delivery of Cas9 Ribonucleoproteins Results in High Levels of Gene Editing in Primary Hepatocytes. CRISPR J 5 (3):397–409.

Romoren, K., X.T. Fjeld, A.B. Poleo, G. Smistad, B.J. Thu et al., 2005 Transfection efficiency and cytotoxicity of cationic liposomes in primary cultures of rainbow trout (Oncorhynchus mykiss) gill cells. Biochim Biophys Acta 1717 (1):50–57.

Schiøtz, B.L., E.G. Rosado, E.S. Baekkevold, M. Lukacs, S. Mjaaland et al., 2011 Enhanced transfection of cell lines from Atlantic salmon through nucoleofection and antibiotic selection. BMC Research Notes 4 (1):136.

Team, R., 2019 RStudio: Integrated Development Environment for R. RStudio, Inc.

Team, R.C., 2020 R: A Language and Environment for Statistical Computing.

Wang, X., L. He, S.M. Goggin, A. Saadat, L. Wang et al., 2018 High-resolution genome-wide functional dissection of transcriptional regulatory regions and nucleotides in human. Nature Communications 9 (1).

Wilberg, R., 2020 Optimization of transfection of primary hepatocytes from Atlantic salmon for functional studies in *Faculty of Chemistry, Biotechnology and Food Science*. Norwegian University of Life Sciences.

Zeilinger, K., N. Freyer, G. Damm, D. Seehofer, and F. Knöspel, 2016 Cell sources for *in vitro* human liver cell culture models. Experimental Biology and Medicine 241 (15): 1684–1698.

