## Supplementary information for "Efficient transfection of Atlantic salmon primary hepatocyte cells for functional assays and gene editing"

**Supplemental data 1:** The Atlantic salmon *elovl5b* promoter region (NC_027327.1: 27244001 – 27245560) cloned into pGL4.10-elovl5bWT.

5’GAGAATGAGGTTAAGGTTAGCAAAAGAGTTAGGGTTAGGCCAAACTAAAAAGCTAAGG
TTGTCAACAACTCTTATCCCCGACGCGACAACTAGATGGCGGTGTACTAGTATCCACAAC
CGCACAGACTATCCCTAAAAAATGCACCAGTATCATAACACAGGTTACTGAGCTCTGCTC
TGTGTAGCCATGTGAGTGATGCAGTGATTTGATTGGTCTCCACTGACGAGAAATGAATGG
GGTGACATCGGCTTGCTAACTGATTTATTAACTGGTTGTGAGTGATGCAGTGATTTGATTG
GTCTCCACTGATGCGAAATGAATGGGGTGACATCGGCTTGCTTACCGACTTGCTATCTGG
GACGCATTCAGCAGTGTTGAAGGTTTTGGAACGTTCAGATAGAAATATGCTAAAATATAG
CTCAAACATGCCTCTCTGGCATATATAAGGAATCACATCGGCTCTAGTCATGGTATTTCTA
TCTGCAACGTTCGAGAACGTTTGGCAACTGAACATGGCCTAGGTTGCAGTTATACACGTA
CCGCGTTCAACCAGTTAGTATGTCGGAAATTCCCGTGAACGCAGCATTGGGCACGTTTTC
TTGTCACTTTCTCTAAATCCAGCATGTCACCTATACTATAGGCTACGAATAAAACCTCACG
AGTAACCTTGTCCCGACTACAACGCACATCCCGGGTGTCTGTAGCCCAGGCAAAATGTAA
GAGTGCACAAATATTATTAAGCACATTGGCTAAAAGAGAATAGCTCTTTGGTATACGTAA
ATGCGCAGTTGTGCATTTAAAACGTGCCTCTACAACTAAGATTTTCACTACTCAGCAAAC
CATGCAGTTTATTAGGCTACAGATTAAATAATGCTAAACTTTACAGGGTGGAGACACTGC
ACGGTGATGAGTTTGATGCTCATTTCCAATACATGATGATCAATGCTTGACCGCTCCATAA
TAATCTCATCATGTAGCTACCCGCACAGCCTACAAGCACTGTAGCTGTTGGCTAGAGCGC
ACATGACAAGACCAGAGTAGACTGGCACATTTGCTATTTAATGCAAGAGATTTTGTCACA
AAACAATCGGTAGAGTTGAAAATGCTATAAACACATTGAATTTAAGAGGAACGGTACAT
GTTGTGTGCACTATGTCATCACGCACTGATTTTTATTTGCAACAAGTCGGTTTGGTGGAAA
AACAATTGGTAGGAAAATGCCCATATTTTCTTTATGCCGGTTTTAAAATATTCACATGAAA
ATCTGTCGCCAATTGAATGGAAACCTAGTTTAAGTGGGACGCCAACATCAGCAGACGGTC
AAATCACCTTCTGGGTGATGATCACTGCCAATGAGGTAGGCCATTAGGGAGGCTGATAGC
CAATTATCTACCATCTGAAGATGATAGCCTAATAGCTAGATCTCGTCACTTATGACGCTCT
GGACAATTTGCCATATAGACTGAGTTCATGTGTTTCTCCCGCTGTTTCCACTGACCGACGA
GGCTGCACATTTGTGCTTTGGGACCTGGGCAGGCAAGATTACGCATCCTCCAGAG3’

**Table S1:** Primers used for cloning pGL4.10-elovl5bWT.

| **Primer name** | **Sequence (5'-3')** |
|---|---|
| El5bWTPromFw | <u>ACTGGCCGGTACCTG</u>GAGAATGAGGTTAAGGTTAGCAA |
| El5bWTPromRv | <u>CCGGATTGCCAAGCT</u>CTCTGGAGGATGCGTAATCTTGC |

*Underlined primer sequences are 15 bp homology arms required for InFusion HD cloning.

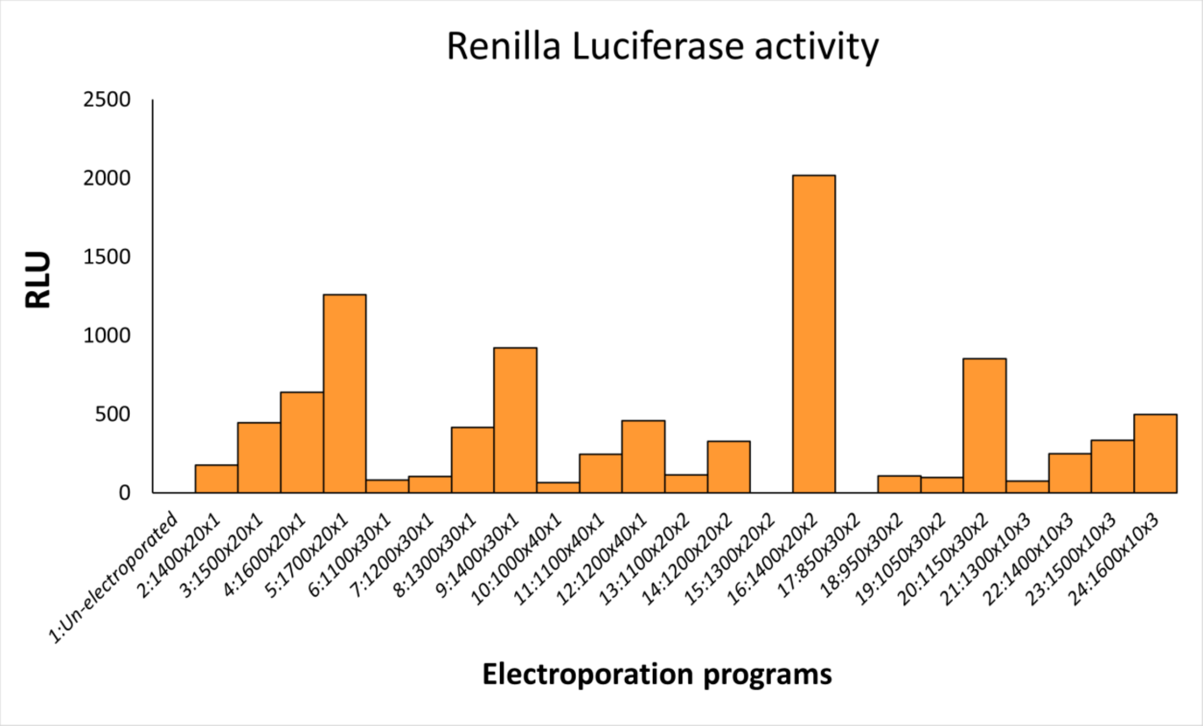


**Figure S1:** Renilla luciferase activity in cells electroporated with 24 different electroporation programs. The Renilla signal is measured in relative light units (RLU). Background signal from un-transfected sample (program 1) is subtracted.
